# A ubiquitin-like protein controls assembly of a bacterial Type VIIb secretion system

**DOI:** 10.1101/2025.01.24.634720

**Authors:** Gabriel U. Oka, Nathanaël Benoit, Axel Siroy, Francesca Gubellini, Esther Marza, Rémi Fronzes

**Affiliations:** CNRS UMR 5234 Microbiologie Fondamentale et Pathogénicité, Institut Européen de Chimie et Biologie, University of Bordeaux, 33600 Pessac, France; Institut Pasteur, Structural Biology of Bacterial Secretion Department, 75015 Paris, France

## Abstract

Type VII secretion systems (T7SS) are crucial bacterial nanomachines that mediate interbacterial competition and host-pathogen interactions in Gram-positive bacteria. Despite their importance, the structural basis for assembly and substrate transport in T7SSb, a widely distributed T7SS variant, remains poorly understood. Here, we present the cryo-EM structure of the T7SSb core complex from *Bacillus subtilis*, revealing how a ubiquitin-like protein, YukD, coordinates assembly of the secretion machinery. YukD forms extensive interactions with the central channel component YukB and promotes its association with the pseudokinase YukC, creating a stable building block for channel assembly. Using microscopy and competition assays, we demonstrate that YukD is essential for proper T7SSb complex formation and contact-dependent bacterial killing. Structural modeling suggests this YukD-dependent assembly mechanism is conserved across diverse Gram-positive bacteria. Our findings reveal how bacteria have adapted a ubiquitin-like protein as a structural regulator for assembling a large secretion complex.

## Introduction

Type VII secretion systems (T7SS) are emerging as important systems for competition and pathogenesis in Gram-positive bacteria, including important human pathogens such as *Mycobacterium tuberculosis, Staphylococcus aureus* and many Streptococcal species^1–3^. By secreting antibacterial toxins or virulence factors, these systems promote bacterial survival by facilitating access to nutrients and ecological niches, directly contributing to virulence during infection and modulating interbacterial competitiveness within biofilms. T7SS were initially classified into two main types - T7SSa found exclusively in diderm Actinobacteria and T7SSb found widely in monoderm Gram-positive bacteria, including many Firmicutes. Recent work has identified eight additional variants (T7SSc-j), expanding the diversity of these systems across many bacterial phyla^4^.

The T7SSa system plays a crucial role in *M. tuberculosis* virulence. Several landmark structural and mechanistic studies have recently shed light on the T7SSa assembly process and its possible modes of substrate secretion^5–9^. The architectural and functional differences between T7SSa and T7SSb are significant, with few homologous core components shared between them. This minimal homology prevents direct extrapolation of structural and functional insights from one system to the other, suggesting that these systems have evolved distinct architectures to serve different biological functions.

The Yuk-Yue system of *Bacillus subtilis* represents a prototypical T7SSb machinery. It comprises the membrane machinery composed of YukB (EssC), YukC (EssB), and YukD (EsaB), along with conserved components such as YueB (EsaA) and YueC (EssA), and secreted WXG100 proteins such as YukE (EsxA)^10^ (Fig. 1A). Throughout the manuscript, we use the *B. subtilis* nomenclature, with the corresponding *S. aureus* protein names provided in parentheses in this paragraph for reference. YukB is a membrane ATPase that contains two N-terminal forkhead-associated (FHA) domains specific to T7SSb, along with a conserved domain of unknown function (DUF) and three C-terminal ATPase domains also found in T7SSa. YukC is a membrane pseudokinase whose full-length transmembrane structure has been determined^11^. Together with YukD, a conserved small ubiquitin-like protein, these proteins are predicted to form a central membrane complex essential for T7SSb function^11^. The T7SSb system contains additional core proteins beyond the central membrane complex. These include: YueB, a multi-spanning membrane protein that features an extended extracellular domain and serves as a receptor for SPP1 phage^12^; YueC, a small membrane protein whose function remains unknown; and YukE, a secreted WXG100 protein that is essential for system function^13,14^.

**Figure 1:**
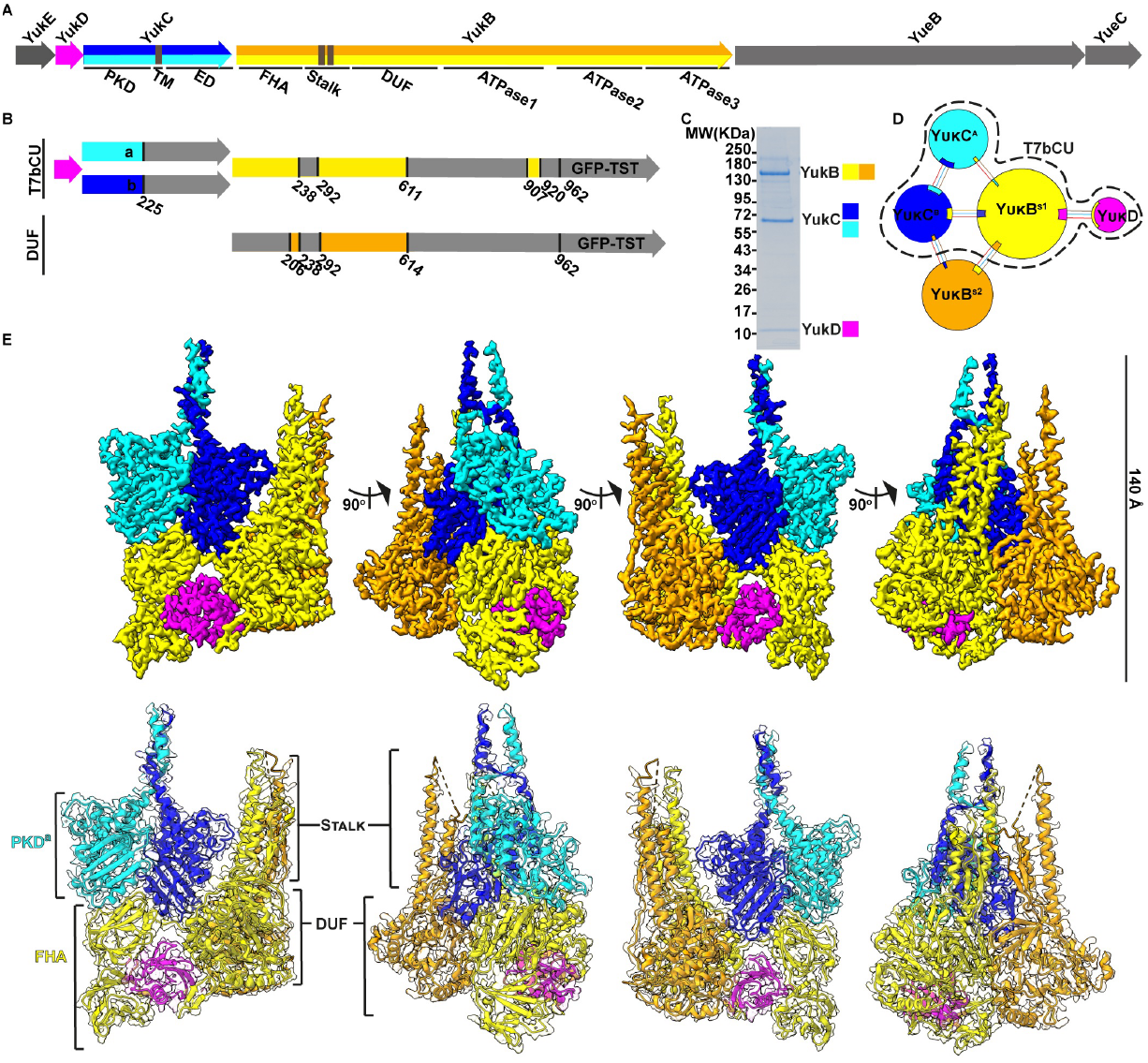
Purification and structural characterization of a T7SSb membrane subcomplex. Organization of the T7SSb operon in *B. subtilis* showing the key functional domains: YukB contains Forkhead-associated (FHA) domains, a stalk region, a domain of unknown function (DUF), and three ATPase domains. YukC comprises a pseudokinase domain (PKD), transmembrane (TM) segments, and an extracellular domain (ED). YukD (magenta) contains a ubiquitin-like domain. YueB and YueC are shown in gray. **(B)** Schematic representation of the protein domains resolved in the CryoEM structure. The T7bCU components are labeled as follows: YukD (magenta); YukC subunit 1 (YukC1a, residues 1-225) and YukC subunit 2 (YukC1b, residues 1-255) in blue and cyan; YukB domains are shown in yellow (residues 3-52, 59-238, and 292-611) and orange (DUF domain of the adjacent YukB subunit, residues 206-238 and 292-614). A C-terminal sfGFP followed by a twin strep tag (GFP-TST) was present but not visible in the CryoEM map. **(C)** SDS-PAGE analysis of the purified T7bCU complex showing YukB(1-962-GFP-TST), YukC, and YukD after affinity chromatography and size-exclusion chromatography purification. **(D)** Schematic repre-sentation of T7bCU assembly showing the 1:2:1 (YukB:YukC:YukD) stoichiometry. The diagram was gen-erated using PDBsum, with component sizes proportional to their accessible surface areas and interface contacts indicated by shared colors. **(E)** Structural analysis of the T7bCU complex. Top: Sharpened Cry-oEM density map (3.8 Å resolution) of T7bCU with an additional YukB DUF and stalk domains. Bottom: Atomic models showing the arrangement of YukB, YukC, and YukD, highlighting the YukB-mediated oli-gomerization interfaces.

In *Bacillus subtilis* and other Gram-positive species, interbacterial competition is mediated by the secretion of proteinaceous effectors from a donor cell into a prey cell^11,15–20^. These effectors belong to a polymorphic family characterized by a conserved N-terminal LXG domain ^18,21^, followed by a non-conserved region and a toxic domain. LXG effectors exhibit diverse toxic activities, including DNase, RNase, and deaminase-like functions, enabling *B. subtilis* to effectively target competing strains. However, the precise mechanisms governing effector selection, recruitment, and delivery by the T7SSb machinery remain poorly understood.

The recruitment and transport of these effectors likely requires specialized regulatory proteins within the T7SSb machinery. One such candidate is YukD, a small ubiquitin-like protein with a structure similar to the C-terminal domain of the T7SSa protein EccD^22^. Recent studies have revealed diverse roles for bacterial ubiquitin-like proteins: they can mediate anti-phage defense through protein conjugation in both Bil and CBASS systems^23–25^ or form higher-order assemblies in response to stress conditions^26^. While the mechanism of action of YukD in the T7SSb system remains to be fully understood, its structural similarity to ubiquitin and its conservation across all but one variant of the T7SS systems^4^ suggest it may play an important regulatory role in the system’s assembly and/or function.

Here, we solved the 3.8 Å CryoEM structure of the *B. subtilis* T7SSb membrane complex, comprising YukB, YukC, and YukD. We show that YukD is a keystone of the Yuk T7SSb membrane core complex that regulates the assembly and the function of the Yuk T7SSb membrane channel.

## Results

### Structural determination of the T7SSb core complex reveals a modular architecture

To elucidate the molecular architecture of the Yuk-Yue T7SSb, we first attempted to purify the native complex directly from *B. subtilis*. A GFP and a strepII-were genomically fused at the Cterminus of YukB in the *B. subtilis* 168 strain. Following bacterial culture, T7SSb complexes were extracted and purified by affinity chromatography. While mass spectrometry analysis of Coomassie-stained bands extracted from SDS-PAGE confirmed YukB as the predominant protein in the purified fraction, the yield and purity levels were insufficient for detailed structural characterization (Supplementary Fig. 1).

To overcome this limitation, we used a heterologous expression system in *E. coli* based on known interactions among the T7SSb components^11^. We constructed a pET-derived plasmid encoding YukD, YukC, and a truncated version of YukB (residues 1-962, named YukBcut). YukBcut contained the N-terminal FHA domain, the stalk comprising the transmembrane segments, the central domain of unknown function (DUF) and the first predicted ATPase domain, all fused with GFP and a Twin-Strep-tag at the C-terminus (Fig. 1B; Supplemental file 1, FASTA). After cell culture, protein extraction by detergents and affinity purification, analysis by SDS-PAGE revealed distinct bands corresponding to YukBcut, YukC, and YukD in the purified complex (Fig.1C).

Cryo-electron microscopy revealed a complex formed by YukBcut, YukC, and YukD. After initial preprocessing of the micrographs, particle picking and 2D classification using CryoSparc^27^, initial *ab initio* 3D reconstructions and consensus 3D refinement revealed an intricate architecture embedded in a detergent belt. Initial analysis of the densities using rigid body docking of atomic models of YukBcut, YukC and YukD obtained using AlphaFold2^28^, allowed the identification of the corresponding densities in this map.

Further analysis using 3D classification and heterogeneous refinements identified three distinct oligomeric states of the complex. Each oligomer contains at least one copy of the same building block comprising YukBcut, YukC and YukD with a 1:2:1 stoichiometry, now referred to as the T7bCU (for Type VIIb Core Unit). Assembly 1 comprises a unique T7bCU in interaction with an additional YukB DUF and stalk domains (Fig. 1, Extended Data Fig. 1A).

Assembly 2 is formed by a dimeric form of T7bCU (Extended Data Fig. 1B), and assembly 3 is a dimer of T7bCU with an additional YukB DUF and stalk domains (Extended Data Fig. 1C). For each assembly, the regions corresponding to the extracellular portion of YukC and the YukBcut ATPase domain were poorly defined. To improve resolution, we performed focused refinement on the T7bCU which is identical across all oligomeric states. We applied a 3D mask encompassing the FHA, stalk and DUF domains of YukBcut, the N-terminal and transmembrane domains of YukC, and YukD, using aligned particles from the consensus refinement as a starting point. Using 303,198 particles, we obtained a 3.8 Å resolution map of the T7bCU and adjacent YukB DUF through focused refinement with a mask encompassing the FHA, stalk and DUF domains of YukBcut, the N-terminal and transmembrane domains of YukC and YukD (Extended Data Fig. 1D and 1E). This map enabled *de novo* atomic building and real-space refinement for the YukB DUF and stalk, with the exception of the transmembrane portion of this domain (residues 249 to 304) as well as the two Nterminal FHA domains (Fig. 2A and 2B). Structure analysis revealed that the structure of the YukB DUF and stalk are similar to that of the corresponding domains in the mycobacterial EccC (PDB: 7NPR) (Extended Data Figure 2A). YukB DUF domain extends toward the N-terminus through a linker (residues 212 to 226) followed by two consecutive FHA domains (Fig. 2A).

**Figure 2:**
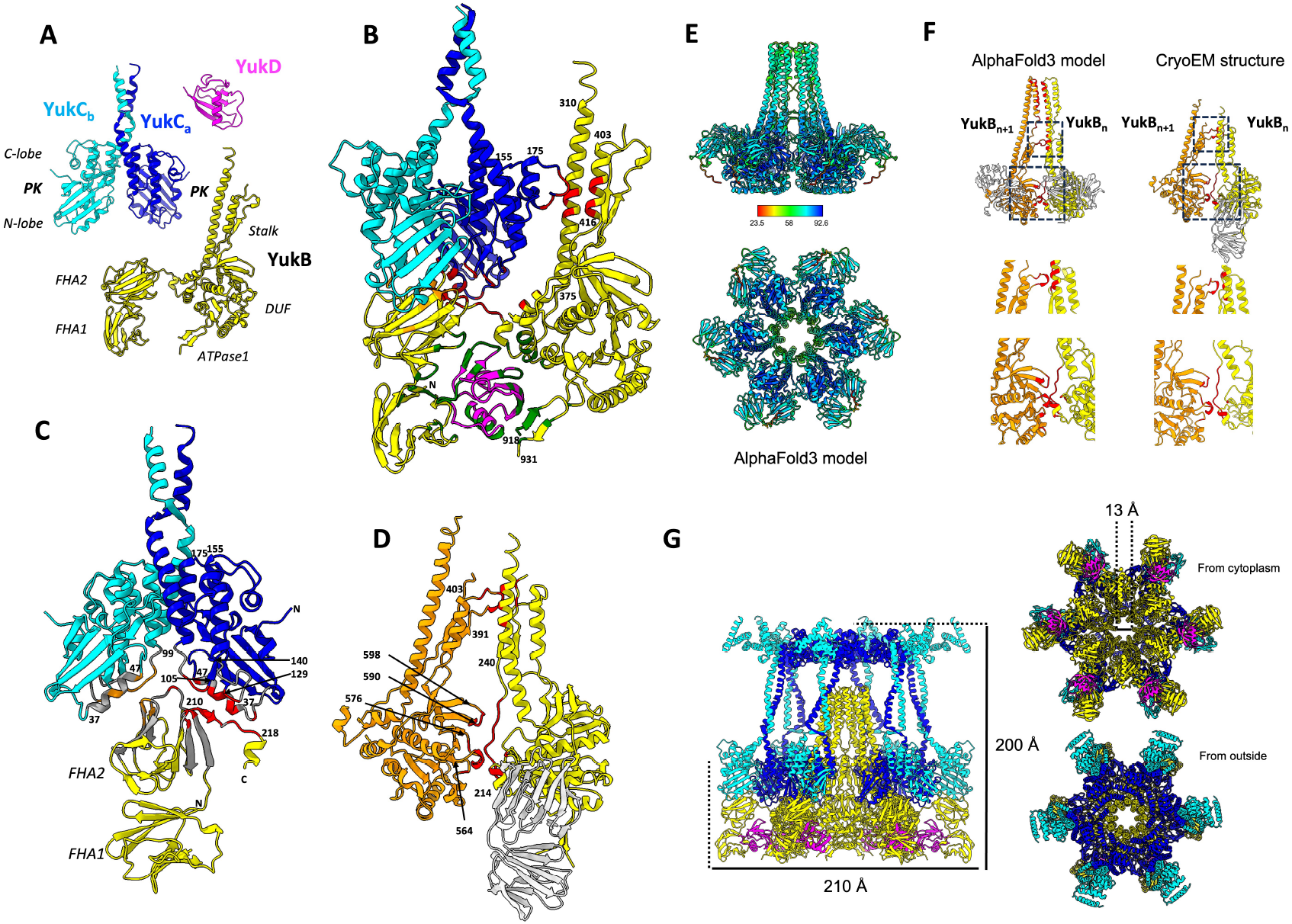
Molecular architecture and interactions within the T7SSb complex. **(A)** Atomic model of the T7bCU showing the structural organization of YukB (yellow), YukC dimer (blue/cyan), and YukD (magenta). The model reveals the arrangement of key domains including YukB’s FHA domains, stalk, and DUF, YukC’s pseudokinase domain and transmembrane regions, and the ubiquitin-like YukD. **(B)** Detailed view of YukD’s extensive interactions with YukB, highlighting in green the interactions between YukD and YukB’s FHA domains, DUF domain, and the beta-hairpin (residues 918-931) from the first ATPase domain. The interactions between YukC and YukB FHA domains, DUF and stalk are highlighted in red.**(C)** Close-up view of the YukC dimer interface with YukB, showing how one YukC subunit (YukCa in dark blue) makes contacts with with YukB through the FHA2 domain and N-terminal linker (residues 210-218) between FHA and stalk domains. The αC-helix (residues 37-47) and connecting loops (residues 99-105 and 129-140) in YukC play key roles in these interactions. **(D)** Analysis of T7bCU oligomerization interfaces, showing in ref the specific contacts between YukB subunits through their DUF and stalk domains. Key interaction regions include the FHA-stalk linker (residues 219-240) and protruding loops from the DUF domain. **(E)** AlphaFold3 model of the YukB hexamer colored by pLDDT confidence score, showing the predicted structural organization of the complex. **(F)** Comparison between the experimental CryoEM structure and the AlphaFold3-predicted YukB hexamer model. The contacts found in the experimental structure and model are shown as zooms below the overall structure. The different position of the FHA is highlighted by a box. **(G)** Complete structural model of the hexameric YukBcut/C/D assembly, showing the central channel formed by YukB components (∼13 Å in diameter) surrounded by six YukC dimers arranged in a crown-like structure.

We were also able to build the pseudo-kinase (PK) domain of YukC (residues 1 to 206) and a portion of the linker just preceding the transmembrane helix (residues 207 to 230) (Fig. 2A and 2B). In our structure, YukC PK domain was found as a dimer (Fig. 2A). Its structure can be superimposed on the previously determined crystal structure (PDB: 6Z0F) (Extended Data Fig. 2B). The main difference between the two structures is the angle of the transmembrane (TM) helices relative to the PK domain (Extended Data Fig. 2B). Although the resolution of the map was insufficient for *de novo* building of the YukC transmembrane helices and C-terminal domains, low-resolution densities enabled rigid-body docking of these regions using the crystal structure as a template (Extended Data Fig. 2B). This revealed that, upon interaction with YukB, YukC undergoes a conformational change with residue 212 as a pivot point. YukD structure was completely resolved (Fig. 2A and 2B) and is found almost identical to the crystal structure determined previously (PDB: 2BPS) (Extended Data Fig. 2C).

Within each T7bCU, YukB acts as a scaffold and interacts both with YukC and YukD. The interaction between YukB and YukD is particularly extensive and represents nearly 30% of YukD’s total surface (1431.5 Å^2^ out of 4921.5 Å^2^) (Fig. 2B). YukB interacts with YukD like a hand catching a ball, with the YukB FHA domains on one side and the DUF domain on the other side. These extensive interactions stabilize the relative conformation of these two parts of YukB (Fig. 2B). Additionally, a small beta-hairpin (residues 918 to 931) located in the first ATPase domain following YukB DUF domain interacts with YukD closing the hand on YukD (Fig.2B). While this loop is welldefined, the rest of the ATPase domain is not visible, indicating high flexibility.

The YukC dimer sits symmetrically onto the tip of the YukB FHA2 domain (Fig. 2C). Both YukC subunits use the same N-terminal lobe regions for these interactions with YukB, with a crucial structural role played by the αChelix (residues 37 to 47).. The αC-helix serves as a molecular switch in protein kinases, but in pseudokinases it has evolved to function primarily as a protein interaction platform (Taylor and Kornev, 2011; Zeqiraj and van Aalten, 2010). Additionally, two loops connecting the N-lobe and C-lobe (residues 99 to 105 and 129 to 140) participate in the interactions with YukB FHA2 in both YukC subunits. Furthermore, one YukC subunit (YukC_a_) makes additional contacts with YukB through the N-terminal part of the linker connecting FHA2 and stalk (residues 210 to 218) (Fig. 2C). A long loop protruding from the C-lobe (Residues 155 to 175) of the YukC_a_ subunit interacts with the side of the YukB stalk region. Additional possible contacts might exist between the loop preceding YukC_a_ αC-helix helix and the DUF domain

Our structural analysis revealed pairs of T7bCUs making specific contacts through their YukB DUF and stalk domains (Fig. 2D). These interactions resulted in assemblies containing up to three YukB subunits, suggesting the formation of a larger complex. For clarity, these three YukB subunits are designated as YukBn, YukBn+1, and YukBn+2, corresponding to their clockwise position when viewing the complex from the cytoplasmic side. The Cterminal part of the linker connecting the FHA and stalk domains (residues 219 to 240) interacts with two loops protruding from the side of the neighboring YukB DUF domain (residues 564 to 576 and 590 to 598). Additional contacts occur between the side of the stalk of the YukB monomer (YukBn) and a loop protruding from the stalk of neighboring YukBn+1 (residues 391 to 403).

We used AlphaFold3 to generate a structural model of the complete oligomeric assembly. Based on the known hexameric organization of homologous ESX inner membrane complexes ^5,8^, we modeled a YukBcut hexamer comprising residues 1 to 603, which encompasses the FHA, DUF and complete stalk and transmembrane domains but excludes the first ATPase domain. The pLDDT scores of the prediction range from 23.6 to 93.6, with most of the predicted structure scoring above 60. The computational model matches our experimental data for the structure and position of the DUF and stalk (RMSD 1.149 Å), while the FHA domains display accurate fold prediction but occupy different positions relative to the DUF (Fig 2E and 2F). Importantly, the YukB-YukB interfaces observed by cryoEM are also present within the predicted YukB hexamer (Fig.2D and 2E. This validation is crucial, as the EccC hexamer lacks contacts between the stalks and DUF domains of neighboring EccC subunits (Extended Data Fig2E) ^8^. The predicted structure of the YukB hexamer differs notably from the EccC experimental structure: the predicted transmembrane helices of YukB align with its stalk domain, unlike in EccB structures where these helices adopt a different orientation relative to the stalk and DUF domains (Extended Data Fig. 2E). The presence of transmembrane helix densities in our unsharpened cryoEM maps supports this predicted orientation of YukB’s transmembrane regions within the detergent micelle.

Using this validated YukB hexamer model as a framework, we built a complete structural model of the YukBcut/C/D hexameric assembly. First, we docked six copies of our T7bCU structure onto the predicted YukB DUFs. We then incorporated the AlphaFold3-predicted transmembrane segments of YukB (residues 239 to 298) to complete regions not resolved in our cryoEM map. Finally, using the YukC crystal structure as a template, we modeled its transmembrane and C-terminal domains (residues 207 to 386), guided by low-resolution densities that indicated their positions.

In the final model, the YukB components (DUF, stalk, and transmembrane segments) assemble to form the central channel of the T7SSb complex (Fig. 2G). Remarkably, this channel of 13 Å in diameter is open, while it is closed in the EccC hexameric structure (Extended Data Fig. 2E). Around this core, six YukC dimers adopt a crown-like structure, with their extracellular domains positioned to form an annular platform at the top through lateral contacts (Fig. 2G). The model suggests additional stabilizing interactions between the N-lobe of each YukC pseudokinase domain and adjacent YukB DUFs (Fig. 2G). The transmembrane regions of YukB and YukC align in a single plane, consistent with their position within the bacterial membrane (Extended Data Fig. 2G).

### YukD stabilizes the assembly of the T7bCUs *in vitro* and *in vivo*

Our structural analysis revealed that YukD plays a crucial role in stabilizing the FHA domains of YukB, as these domains are not visible in cryo-EM maps lacking YukD due to their increased flexibility (Figure 1 and Extended Data Figure 1). To investigate how YukD affects T7bCU assembly and stability, we conducted both *in vitro* and *in vivo* analyses.

For our *in vitro* studies, we generated, purified and compared three variants: the native T7bCU, a YukD-deletion variant (T7bCUs Δ*yukD*), and a YukC-deletion variant (T7bCUs Δ*yukC*). These complexes were purified using affinity chromatography using the TwinStrep tag fused to YukBcut followed by size-exclusion chromatography. For SDS-PAGE analysis, we loaded equal amounts of each sample based on the YukB-GFP band intensity to ensure comparable YukB quantities across all variants. SDS-PAGE analysis revealed that the absence of YukD significantly reduced YukC association, while YukD yield remained unaffected by YukC deletion (Figure 3A). Quantitative analysis confirmed these observations: the YukC/YukB ratio was markedly lower in T7bCUsΔYukD compared to native T7bCU, while the YukD/YukC ratio remained consistent between native T7bCU and T7bCUsΔYukC (Figures 3B and 3C).

**Figure 3:**
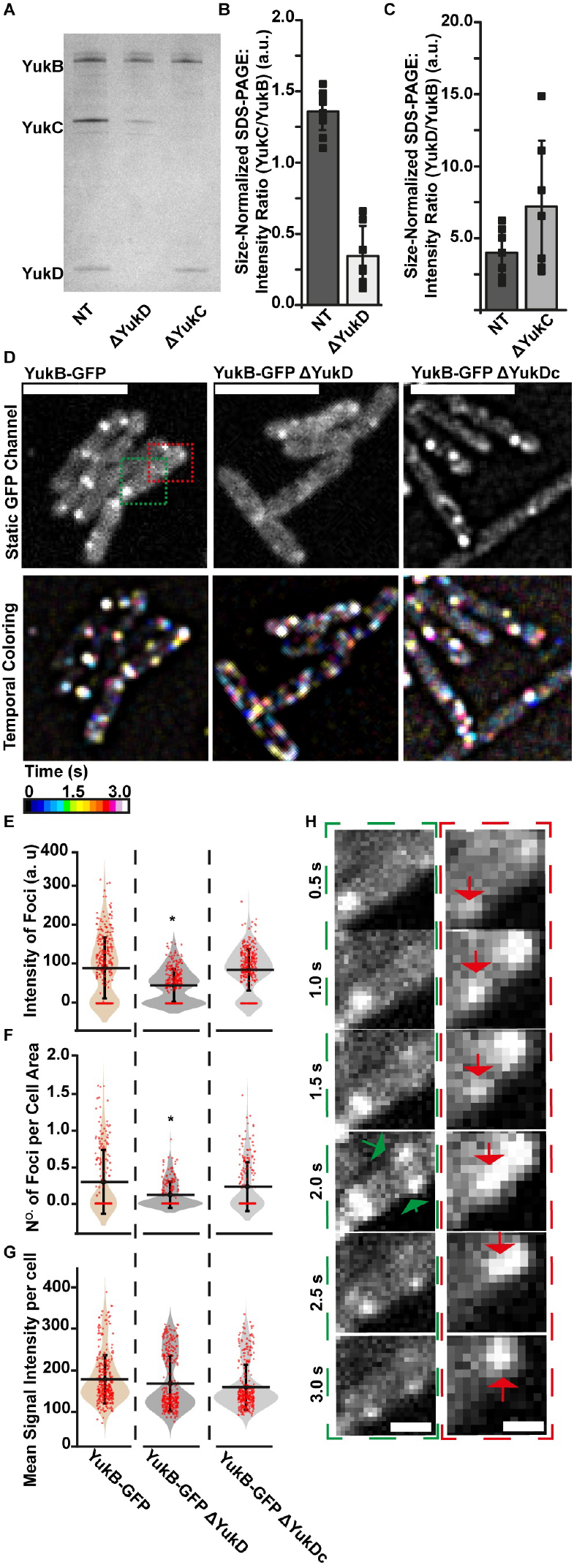
YukD is Essential for T7SSb Complex Assembly. SDS-PAGE analysis comparing purified native T7bCU (NT), YukD-deletion (ΔYukD), and YukC-deletion (ΔYukC) variants after size-exclusion chromatography. **(B)** Quantification of YukC/YukB ratios from SDS-PAGE for NT and ΔYukD samples. One-way ANOVA shows significant difference (F(1, 17) = 168.93, P < 0.0001). **(C)** Quantification of YukD/YukB ratios from SDS-PAGE for NT and ΔYukC samples. No significant difference observed (One-way ANOVA: F(1, 12) = 2.918, P = 0.113). Data points represent independent purification fractions (n=2). Mean ± SD shown. **(D)** Live-cell microscopy of *B. subtilis* strains. Top: YukB-GFP localization in wild-type, ΔYukD, and complemented (ΔYukD + YukD) strains. Bottom: Temporal color-coding showing dynamic behavior of YukB-GFP foci. Scale bars: 5 µm. **(E-G)** Quantitative analysis of YukB-GFP localization. Cells were grown in parallel under identical conditions, imaged on CM agarose pads within 1-2 hours, and analyzed using MicrobeJ with consistent parameters. **(E)** Foci intensity distribution (n=330 per strain). ANOVA F(2, 987) = 56.22, P < 0.0001, with Tukey’s test showing significant differences between WT vs ΔYukD (P < 0.0001) and ΔYukD vs ΔYukDc (P < 0.0001). Asterisk indicates significant difference from WT.**(F)** Foci density per cell area (n=200 cells). ANOVA F(2, 597) = 14.32, P < 0.0001, with significant differences between WT vs ΔYukD (P < 0.0001) and WT vs ΔYukDc (P = 0.002). **(H)** High-speed imaging (500 ms/frame) of YukB-GFP dynamics. Red arrows: movement along cell envelope; green arrows: foci appearance/repositioning. Scale bar: 1 *µ*m.

To examine the role of YukD *in vivo*, we performed fluorescence microscopy using three *B. subtilis* strains: the parental strain carrying a genomic *yukB-gfp* fusion (YukB-GFP), its derivative lacking *yukD* generated by recombination and subsequent excision of a kanamycin resistance cassette (YukB-GFP ΔYukD), and a complemented strain where *yukD* was reintroduced ectopically (YukB-GFP ΔYukD + YukD) (see material and methods section for details). The parental strain exhibited well-defined, randomly distributed YukB-GFP foci throughout the cell, consistent with T7SSb complex formation. In contrast, the Δ*yukD* strain showed diffuse, less intense foci, suggesting impaired complex assembly (Figure 3D and movies S1-3). Quantitative analysis demonstrated significant reductions in both foci intensity (Figure 3E) and foci density per cell area (Figure 3F) in the Δ*yukD* strain compared to the parental strain of *B. subtilis* expression the YukB. Importantly, complementation with ectopically expressed YukD restored foci formation to levels comparable to the parental *B. subtilis* YukB-GFP strain (Figures 3D-F). Temporal coloring of time-lapse fluorescence images further revealed active movement, assembly, and redistribution of YukB-GFP in the parental and complemented strains, which were markedly reduced and dispersed in the ΔyukD mutant (Figure 3D, 3H). Within a 3-second time span, we observed YukB-GFP foci traveling distances of 1–2 micrometers, either parallel to the cell poles or in crossed directions, highlighting that some foci exhibit high dynamic behavior (Figure 3D, 3H). However, the majority of YukB-GFP foci observed in the parental and complemented strains demonstrate limited dynamics, remaining confined within submicrometer spatial regions (Figure 3D, 3H). Total cellular fluorescence intensity remained constant across all strains (Figure 3G), confirming that the differences in foci formation reflected YukD’s contribution to complex stability rather than variations in YukB-GFP expression levels.

### YukD is required for interbacterial killing via T7SSb

To investigate the role of YukD in T7SSb-mediated bacterial competition, we performed both endpoint colony competition assays and real-time microscopy analysis.

For colony competition assays, we used kanamycin-resistant donor strains (wild-type *B. subtilis* 168, ΔyukB, YukB-GFP, and YukB-GFP ΔyukD) competing against a spectinomycin-resistant ΔyukB recipient strain expressing RFP. The recipient strain also lacked the yxiD-yxxD immunity operon^11,18,29^. After 36 hours of co-culture on CM agarose plates, we quantified colony-forming units (CFUs) to determine survival ratios (Fig. 4A). The wild-type strain showed robust competitive ability with a three-log advantage in CFU ratio, while the YukB-GFP strain maintained a two-log advantage. In contrast, both ΔyukB and ΔyukD strains showed no competitive advantage, with CFU ratios approaching 1, indicating complete loss of T7SSb killing activity.

**Figure 4:**
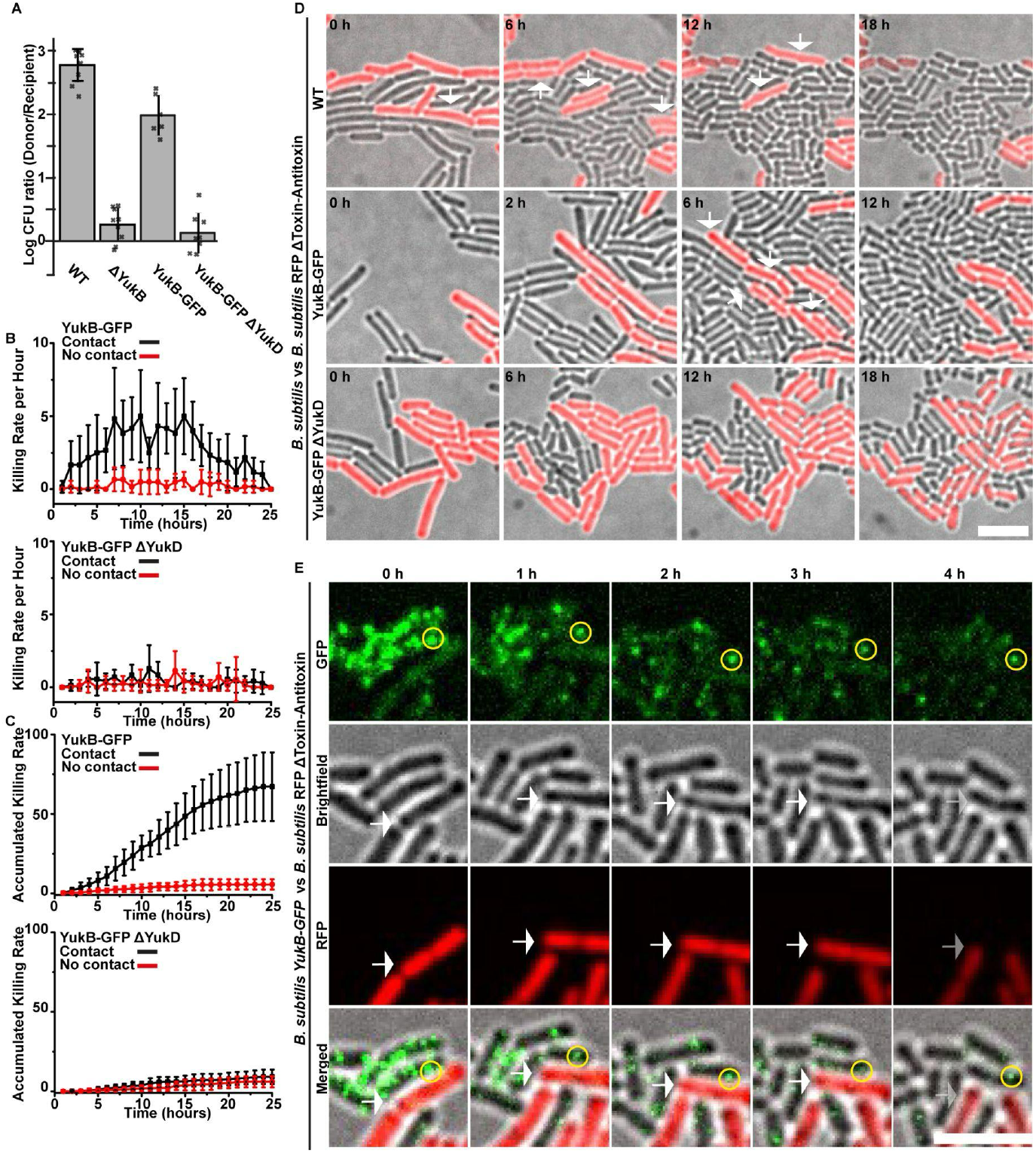
YukD is Required for T7SSb-Mediated Bacterial Competition. **(A)** Quantification of bacterial competition assays. Donor (*B. subtilis* WT, ΔYukB, YukB-GFP, YukB-GFP ΔYukD) versus recipient (ΔYukB-RFP) strains were co-cultured for 36h on CM agarose. Donor/recipient ratios calculated from CFU counts. Error bars: SD from 3-4 independent experiments, each point representing mean of duplicate measurements. **(B)** Killing rate analysis by time-lapse microscopy. Contact-dependent (black) and contact-independent (red) killing events quantified per hour. Contact-dependent: fluorescence loss at donor-recipient contact points; contact-independent: loss without visible contact. Data from six 132µm × 132µm fields per strain. Images captured every 15min (brightfield) and 1h (fluorescence) over 24h (Movies S4-6). Error bars: SD. **(C)** Cumulative killing events over time, categorized as contact-dependent (black) or contact-independent (red) as described in (B). **(D)** Representative time-lapse images showing donor cells (brightfield) killing recipient cells (RFP). White arrows indicate RFP signal loss and cell envelope disappearance. Merged images show brightfield and RFP channel overlay. **(E)** T7SSb localization during killing (0-4h). Top: YukB-GFP dynamics in donor cells. Bottom: Merged brightfield/RFP showing recipient cell death. Yellow circles track GFP foci associated with target cell death. Scale bar : 5 *µ*m.

So far, these results demonstrate that a functional T7SS provides a significant adaptive advantage in interbacterial competition, consistent with its role as described in our previous work and other studies ^11,18^. However, our findings reveal a critical and previously uncharacterized role for YukD in mediating this advantage. To obtain more robust evidence of direct killing mediated by the T7SS, we performed time-lapse fluorescence microscopy of the competition (Fig. 4B-E, Supplementary Movies X-Y). We tracked killing events by monitoring the loss of RFP fluorescence in recipient cells, quantifying both the killing rate (Fig. 4B) and cumulative killing over time (Fig. 4C). Killing events were characterized by sudden loss of prey cell fluorescence followed by cell envelope disappearance (Fig. 4D). The YukB-GFP strain showed peak killing activity between 10-18 hours, primarily during direct contact with prey cells. In contrast, the ΔyukD strain showed negligible killing rates, with no distinction between contact-dependent and contact-independent cell death events (Figs. 4B, 4C). Supporting the contact-dependent nature of killing, we observed that T7SSb foci in attacker cells frequently localized near prey cells prior to their death, maintaining this position throughout the killing process (Fig. 4E).

## Discussion

### Novel Structural Insights Reveal a Conserved Architecture of T7SSb Systems Across Gram-positive Bacteria

In this study, we present the first high-resolution structure of a T7SSb subcomplex, providing insights into the architecture and assembly of this bacterial secretion system. This structural data, combined with computational predictions, enabled us to generate the first detailed model of the T7SSb membrane core channel formed by YukB, YukC, and YukD. Comparison with T7SSa structures reveals significant structural differences that are in line with the specialized roles of these secretion systems. While the central YukB component shares structural homology with its T7SSa counterpart EccC in its DUF and stalk domains, YukB uniquely possesses a N-terminal FHA domain that is absent in T7SSa systems. This FHA domain plays a crucial role in T7SSb-specific interactions, particularly in mediating the association with YukC and YukD. Notably, the linker region between the stalk and FHA domains (residues 210-240) serves a dual function, participating in both YukC binding and YukB oligomerization. The overall architecture of the T7SSb complex differs notably from its T7SSa counterpart. In T7SSa, the EccD protein forms a ring around the EccC transmembrane regions and maintains EccC subunits in an ‘open’ conformation through extensive cytoplasmic contacts with their DUF domains, preventing lateral interactions between neighboring EccC subunits at both DUF and stalk levels. In contrast, our T7SSb structure reveals that YukB subunits engage in direct lateral contacts through both their DUF domains and stalk regions, forming a more compact arrangement. This organization is stabilized by the dual architectural roles of YukC: its dimers form a cytoplasmic scaffold that maintains YukB oligomer in its compact conformation, while also organizing into an extracellular ring that could serve as a specialized interface. The T7SSb complex we observe displays an open channel configuration of approximately 13 Å in diameter, while available T7SSa structures show a closed state (Fig 2G and Extended Data Fig. 2). These different conformations might either represent distinct functional states of these secretion systems or fundamental architectural differences, potentially reflecting different stages in the substrate transport cycle. Importantly, structural modeling of T7SSb complexes from diverse Gram-positive bacteria, including *Listeria monocytogenes, Clostridium acetobutylicum*, and *Lactococcus cremoris*, reveals that this core architecture is highly conserved (Fig. 5A and B). While these species show variations in accessory components that likely reflect adaptation to different ecological niches (Fig. 5B), the fundamental organization of the YukB/C/D complex appears to be maintained (Fig. 5A), suggesting its essential role in T7SSb function across Gram-positive bacteria.

**Figure 5:**
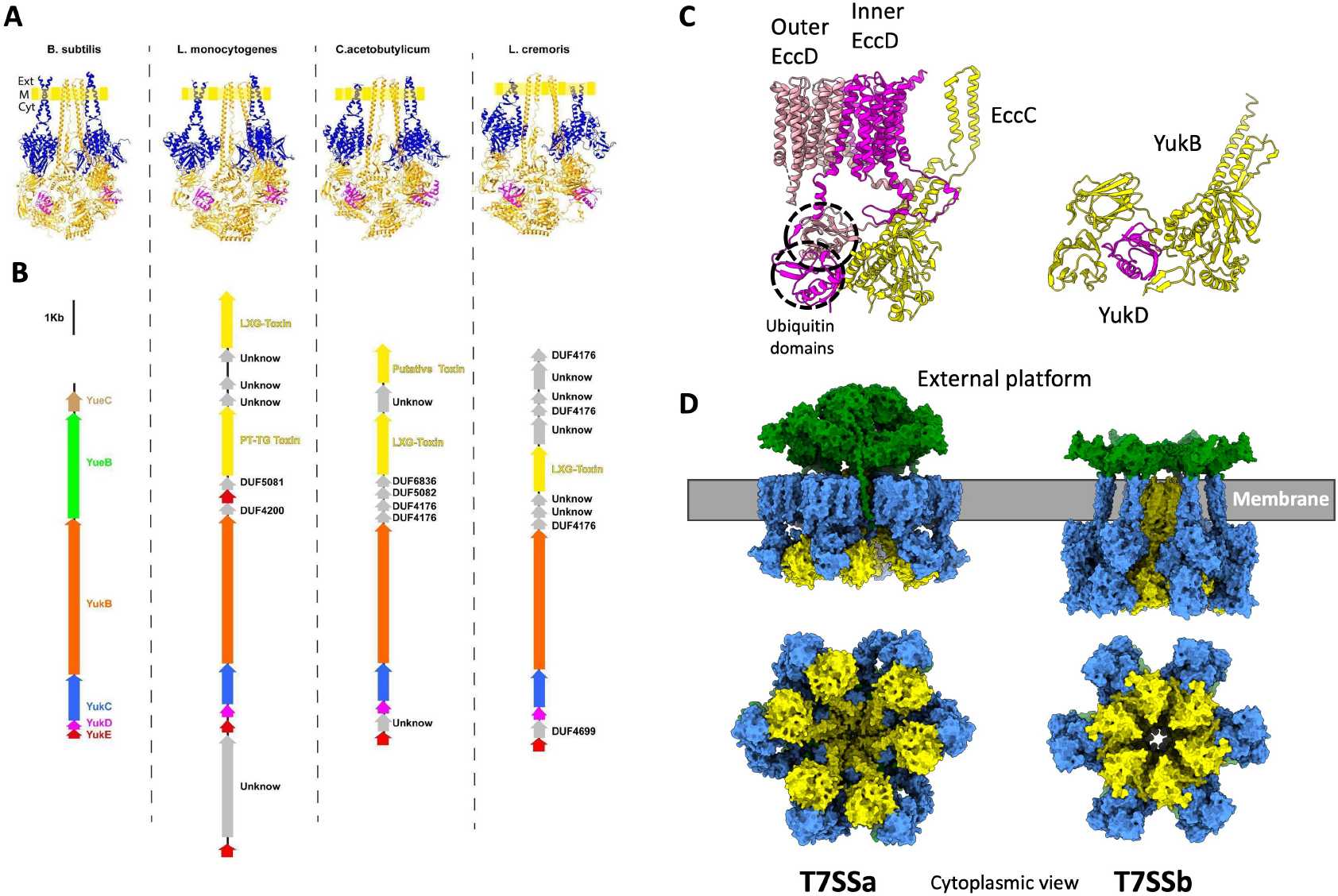
Conservation and comparative analysis of T7SSb and T7SSa architectures. **(A)** Structural models of the T7SSb complexes from diverse Gram-positive bacteria (*Listeria monocytogenes, Clostridium acetobutylicum*, and *Lactococcus cremoris*) predicted by AlphaFold, showing conservation of the core YukB/C/D architecture including the beta-hairpin interaction between YukD and the first ATPase domain. **(B)** Comparative analysis of T7SSb genetic organization across bacterial species, highlighting the conserved core components (YukB, YukC, and YukD) and variable accessory components that reflect adaptation to different ecological niches. **(C)** Comparison of ubiquitin-like domain interactions with EccC and YukB. In T7SSa, two EccD subunits use their N-terminal ubiquitin domains to interact with different regions of the EccC DUF domain, while in T7SSb, a single YukD makes extensive contacts with the YukB DUF domain. **(D)** Comparison of overall organization between T7SSa and T7SSb systems. T7SSa forms a periplasmic platform with a dome-like chamber composed of three EccB dimers capped by MycP proteases (in green), while T7SSb organizes YukC dimers into an extracellular crown-like structure (green). The DUF and stalk domains of EccC and YukB (in Yellow) are found in an open and closed conformations while the transmembrane channels are closed and open respectively in the T7SSa and T7SSb complexes. EccD and YukC associated to the YukB FHA and YukD form peripheral scaffold that maintain EccC and YukB in their respective conformation.

### YukD: A Ubiquitin-like Protein with a Central Role in T7SSb Assembly and Function

The near-universal conservation of YukD across T7SSb variants^9,19^, with only one known exception, underscores its fundamental importance to these secretion systems. The essential nature of YukD is demonstrated by our *in vivo* studies, where its deletion results in diffuse, poorly organized T7SSb complexes rather than the well-defined foci observed in wild-type cells, and abolishes T7SSb-mediated bacterial competition. These findings position YukD as a central architectural element whose presence is required for both the assembly and function of the T7SSb system. We show that YukD makes extensive interactions with YukB that are critical for membrane complex stability, engaging with its FHA domains and promoting their interaction with YukC’s pseudokinase domain. In T7SSa, EccD assembles as a dimer where both subunits interact with EccC through their N-terminal ubiquitin domains (Fig. 5C). The inner EccD subunit’s ubiquitin domain binds to the same region of the EccC DUF that YukD contacts in T7SSb, while the outer EccD subunit’s ubiquitin domain interacts with a different region of the DUF, together providing a dual anchor point for the EccD-EccC interaction (Fig. 5C). YukD also interacts with a beta-hairpin (residues 918-931) protruding from the first ATPase domain. Notably, this interaction appears to be a conserved feature, as structural modeling predicts similar interactions in T7SSb systems across diverse Gram-positive species (Fig. 5A). The conservation of this interaction across T7SSb species suggests it may play an important role in coupling YukD-mediated complex assembly to ATPase activity and T7SSb function. While AlphaFold predictions suggest that the T7SSa protein EccC5 contains a similar extension within its first ATPase domain, the confidence of the structural prediction of its interaction with the N-terminal ubiquitin domain of EccD is very low (Supplementary Figure 3). Nevertheless, the presence of this extension in both systems raises the possibility of a shared architectural feature. The structural role we have uncovered for YukD represents a distinctive adaptation of a ubiquitin-like fold. While eukaryotic ubiquitin-like proteins typically function through reversible conjugation to target proteins^30,31^, YukD acts as a permanent structural component that stabilizes complex assembly through non-covalent interactions. This represents a novel use of the ubiquitin fold beyond its well-characterized roles in protein modification^32^ and sulfur-transfer pathways^31^, demonstrating how bacteria have adapted this ancient protein fold for structural roles in large molecular machines.

### Mechanisms of T7SSb Assembly and Its Role in Contact-Dependent Killing

Our structural and functional analyses reveal how T7SSb assembly is coordinated. The key positioning of YukD in the complex, bridging the FHA domain and ATPase region of YukB while promoting interaction with YukC, suggests it may serve as a molecular switch coordinating complex assembly with activation. In addition, while T7SSa and T7SSb have evolved distinct architectures, they both organize components in the periplasmic/extracellular space. In T7SSa, EccB and MycP form a defined periplasmic platform that includes a dome-like chamber formed by three EccB dimers capped by a trimer of MycP proteases (Fig. 5D). In T7SSb, the C-terminal domains of YukC dimers form a crown-like structure and organize into an extracellular ring (Fig. 5D). This extracellular interface could be further completed by other T7SSb components not present in our structure, such as YueB, which contains an extensive extracellular domain, YueC, or other yet unidentified components. The presence of organized extracellular structures across these systems, despite their different compositions, suggests their importance for substrate processing or targeting.

Our microscopy studies reveal that properly assembled T7SSb complexes exhibit distinct spatial and temporal organization within bacterial cells. In wild-type cells, these complexes form well-defined, dynamic foci. This mobility appears to be functionally significant, as we observe T7SSb foci specifically localizing to points of contact with target cells prior to killing events. The correlation between foci positioning and subsequent cell death suggests that the T7SSb complex must be fully assembled and properly localized for efficient substrate delivery, with the frequency of killing increasing after several hours of interbacterial competition. Furthermore, the T7SSb-mediated killing differs significantly from T6SS, T4SS, and Tad systems by operating through a slower, sustained-contact mechanism. The observed contact dependence of T7SSb-mediated killing raises intriguing questions about substrate recognition and transport mechanisms. The extracellular platform formed by YukC dimers could serve as a specialized interface for target cell recognition or substrate processing. Understanding how complex assembly and activation are coordinated with substrate recognition and delivery represents an important direction for future investigation.

## Acknowledgements

We thank the Bordeaux Imaging Center for access to the fluorescence microscopes and the IECB for access to the electron microscopes. We thank Dr. Kazuo Kobayashi for discussions and Dr. Daniel B. Kearns for providing *B. subtilis* DK1042 comIQ12L strain.

## Materials and Methods

### Bacterial Strains, Growth Conditions, and Media for Experimental Assays

Oligonucleotides, plasmids and bacterial strains utilized in this study are described in Tables S2, S3 and S4. Briefly, the antibiotics utilized included ampicillin (100 μg/mL), kanamycin (10 μg/mL), and spectinomycin (100 μg/mL). *Escherichia coli* strains BL21-AI (Invitrogen) and DH5α were typically cultured in Luria-Bertani (LB) broth or 1% agar supplemented with ampicillin when carrying the plasmids pGUO19.1WT, pGUO19.1KOYukD, and pGUO19.1KOYukC. *B. subtilis* strains were grown in LB or MC as described in ^11^. Interbacterial competition assays were performed on competitive medium (CM).

### Cloning and Gene-Knockouts

Target genes were amplified using PrimeSTAR Max DNA Polymerase (Takara) with genomic DNA from *Bacillus subtilis* 168 ^33^ or synthetic DNA (Genscript) as the template. The oligonucleotides used for amplification are listed in Table S1. The pGUO19.1WT plasmid was used as a template for PCR amplification with pairs of oligos (Table S1) flanking the YukD and YukC genes for knockout. PCR products were treated with DpnI, purified, and incubated at 50°C for 1 hour before transformation into E. coli DH5α via heat shock. Isolated colonies grown on LB agar supplemented with ampicillin were selected, plasmids were extracted, and DNA sequencing was performed to confirm the gene deletions. For the engineering of *B. subtilis*, steps were performed as previously described as in^11^. Briefly, 5 mL of culture was grown in MC medium for five hours at 37°C. Subsequently, 5 µL (∼300 ng) of synthetic plasmids (Table S3) containing a kanamycin resistance marker flanked by lox sites and 500-base regions for allelic recombination at the 5’ and 3’ extremities were added to the culture, followed by incubation for an additional 2 hours at 37°C and 180 rpm. The cells were harvested and plated on LB agar supplemented with kanamycin at 37°C. Isolated colonies were then selected and subjected to a second round of transformation with pDR244. Transformants were selected on LB agar plates containing spectinomycin and grown at 30°C for 12 hours. Colonies able to grow on spectinomycin but susceptible to kanamycin were selected, and correct clones were verified by PCR and genomic DNA sequencing.

### Expression and Purification

*E.coli BL21-AI™ One Shot* (Invitrogen) colonies transformed with plasmids pGUO19.1WT and derivatives were typically inoculated into 5 mL of LB broth medium and grown overnight at 37°C with shaking at 200 rpm. The cultures were then diluted into 4 L of LB medium and grown at 37°C until an optical density of 0.7 at 600 nm was reached. Induction was carried out by adding 0.5 mM IPTG and 0.1% arabinose, followed by incubation at 18°C for 16 hours with shaking at 200 rpm. Cells were then harvested by centrifugation, resuspended in a lysis buffer (50 mM Tris-HCl, pH 8.0, 300 mM NaCl), and lysed using an Emulsiflex. The lysates were subjected to centrifugation for cell clarification (30,000g for 30 minutes) followed by membrane pelletization (180,000g for 45 minutes). The membrane pellet was resuspended in buffer A (50 mM Tris-HCl, pH 8.0, 300 mM NaCl, 0.5% n-Dodecyl β-D-maltoside (DDM) (Anatrace)) and homogenized using Potter-Elvehjem homogenizers before undergoing a secondary clarification step (30,000g for 30 minutes). Purification was performed using FPLC (ÄKTA avant system) at a flow rate of 5 mL/min with a 5 mL StrepTrap™ XT prepacked chromatography column (Cytiva). Eluted fractions were subsequently subjected to size exclusion chromatography using a Superose™ 6 Increase column (Cytiva). All steps were conducted at 5°C and completed on the same day. Pure proteins were identified by SDS-PAGE, and images were acquired using the ChemiDoc™ System (Bio-Rad). Quantitative analysis was performed using Image Studio™ Software (LI-COR). The purity was confirmed by SDS-PAGE and Western Blot analysis. Color Prestained Protein Standard (NEB: P7719S) was employed for SDS-PAGE analysis.

### Cryo-EM sample preparation and data acquisition

Freshly eluted samples were typically applied onto R2/2 Cu 200 mesh grids (Quantifoil) that had been glow-discharged for 45 seconds at 2.0 mA using a Vitrobot Mark IV (Thermo Fisher Scientific), before being plunge-frozen in liquid ethane. Cryo-images of the purified samples were recorded with a Talos Arctica microscope (Thermo Fisher Scientific) operated at 200 kV and equipped with a K2 Summit direct electron detector (Gatan). Dose-fractionated data were collected within a defocus range of 0.2 to −2.0 µm at 36,000x magnification, corresponding to a pixel size of 1.2 Å, using SerialEM. Three datasets were collected (table 1) with a total electron dose of approximately 50 electrons/Å^2^.

**Table 1.** Cryo-EM data collection and refinement statistics.

|  |  |  |
| --- | --- | --- |
| Table S1- |  |  |
| Model |  |  |
| Data collection | Talos Arctica (Thermo Fischer) |  |
| Voltage (kV) | 200 |  |
| Camera | K2 Summit (Gatan) |  |
| Magnification | 36,000 X |  |
| Electron dose per frame (e- /Å<sup>2</sup> ) | 50 |  |
| Pixel size (Å) | 1.2 |  |
| Defocus range (µm) | -0.2 /-2.5 |  |
| Processing | Single Particle Analysis |  |
| Symmetry imposed | none |  |
| Micrographs (no.) | 15820 |  |
| Initial particle images (no.) | 6,311,712 |  |
| Final particle images (no.) | 303,198 |  |
| Map resolution (Å) | 3.77 |  |
| FSC threshold model | 0.143 |  |
| ===== |  |  |
| Composition |  |  |
| Chains | 5 |  |
| Atoms | 11945 (Hydrogens: 0) |  |
| Residues | Protein: 1441 Nucleotide: 0 |  |
| Water | 0 |  |
| Ligands | 0 |  |
| Bonds (RMSD) |  |  |
| Length ( Å ) ( # > 4σ ) | 0.004 (0) |  |
| Angles ( ° ) ( # > 4σ ) | 0.950 (0) |  |
| MolProbity score | 1.54 |  |
| Clash score | 4.26 |  |
| Ramachandran plot (%) |  |  |
| Outliers | 0.07 |  |
| Allowed | 4.71 |  |
| Favored | 95.22 |  |
| Rama-Z (Ramachandran plot Z-score, RMSD) |  |  |
| whole ( N = 1423 ) | -0.82 (0.22) |  |
| helix ( N = 465 ) | 0.33 (0.25) |  |
| sheet ( N = 246 ) | -1.03 (0.32) |  |
| loop ( N = 712 ) | -0.89 (0.23) |  |
| Rotamer outliers (%) | 0.99 |  |
| Cβ outliers (%) | NA |  |
| Peptide plane (%) |  |  |
| Cis proline/general | 1.9/0.0 |  |
| Twisted proline/general | 0.0/0.0 |  |
| CaBLAM outliers (%) | 2.06 |  |
| ADP (B-factors) |  |  |
| Iso/Aniso ( # ) | 11945/0 |  |
| min/max/mean |  |  |
| Protein | 38.66/177.84/73.97 |  |
| Nucleotide | --- |  |
| Ligand | --- |  |
| Water | --- |  |
| Occupancy |  |  |
| Mean | 1.00 |  |
| occ = 1 (%) | 100.00 |  |
| 0 < occ < 1 (%) | 0.00 |  |
| occ > 1 (%) | 0.00 |  |
| Data |  |  |
| ===== |  |  |
| Box |  |  |
| Lengths ( Å ) | 110.40, 133.20, 151.20 |  |
| Angles ( ° ) | 90.00, 90.00, 90.00 |  |
| Supplied Resolution ( Å ) | 3.9 |  |
| Resolution Estimates ( Å ) | Masked | Unmasked |
| d FSC (half maps; 0.143) | 3.7 | 3.7 |
| d 99 (full/half1/half2) | 4.6/2.5/4.6 | 4.6/2.5/4.6 |
| d model | 4.1 | 4.2 |
| d FSC model (0/0.143/0.5) | 3.6/3.8/4.0 | 3.8/3.8/4.1 |
| Map min/max/mean | -0.25/0.60/0.02 |  |
| Model vs. Data |  |  |
| ===== |  |  |
| CC (mask) | 0.83 |  |
| CC (box) | 0.76 |  |
| CC (peaks) | 0.73 |  |
| CC (volume) | 0.83 |  |
| Mean CC for ligands | --- |  |

### CryoEM Image Processing

All data processing was conducted using CryoSPARC^27^, and the cryo-EM data processing workflow is schematized in Fig. S2. Briefly, movies were aligned to correct for beam-induced motion using the “Patch Motion Correction” function, and Contrast Transfer Function (CTF) parameters were estimated with “Patch CTF Estimation.” Particles were initially picked using the Template Picker, with an AlphaFold-based model serving as the initial template. These particles underwent 2D classification, *Ab initio* modeling, and heterogeneous refinement. To improve particle quality, Topaz training and Topaz particle extraction were employed, followed by additional cycles of 2D classification, non-uniform (NU) refinement, heterogeneous refinement and local refinement to further refine the maps. Final heterogeneous refinement and 3D reconstruction were performed to identify particles corresponding to assembly 1, assembly 2, and assembly 3.

### Building and Refinement of the Atomic Model

The model of the T7CP, generated using AlphaFold ^28,34,35^, was docked into the refined cryo-EM map using the “Fit in Map” function in ChimeraX ^36^. The map was sharpened using DeepEMhancer ^37^, and the final model underwent several rounds of manual refinement in Coot ^38^ and ISOLDE ^39^, followed by automated real-space refinement with phenix real space refine. Details of the cryo-EM refinement statistics and the FSC map versus model plot are provided in Table 1, respectively. The final model was validated using MolProbity^40^ and phenix.validation_cryoem, as implemented in the PHENIX software suite^41^.

### Time-Lapse Fluorescence Microscopy

After cultivation in LB broth, *B. subtilis* DK and 168 cells were collected by centrifugation (6,000 × g for 5 minutes), washed twice in LB broth, and normalized to an OD_600_ of 2.0. Cells were pipetted onto MC1x-agarose supports and observed using an inverted DMI8 microscope (Leica) equipped with a Confocal Scanner Unit CSU-W1 T2 (Yokogawa), a Prime 95B sCMOS camera (Photometrics), and an HCX PL Apo 100× oil immersion objective (NA 1.46). Fluorescent images for foci analysis were acquired by exciting GFP-and RFP-tagged cells using laser diodes at 488 nm and 561 nm, respectively. Exposure times ranged from 50 ms to 500 ms per frame, with images streamed over a duration of seven seconds. Image processing was performed using FIJI (Schindelin et al., 2012), with background subtraction and signal summation over two seconds of excitation in the GFP channel. To monitor interbacterial competition, brightfield images were captured every 15 minutes, while fluorescence images (GFP and RFP) were acquired every 60 minutes over a 24-hour period at room temperature. All image processing and analysis were performed manually using FIJI software ^42^ and MicrobeJ software ^43^.

### Interbacterial competition assays

For the bacterial competition assay based on viability, *Bacillus subtilis* strains were used. The prey strain carried pDR244, which confers resistance to spectinomycin (100 µg/mL), while the attacker strain carried pBE-S, which confers resistance to kanamycin (10 µg/mL). Both strains were grown overnight in LB medium at 37°C as pre-inocula, with the respective antibiotics. Subsequently, the cultures were diluted 1:50 in fresh LB medium supplemented with the appropriate antibiotics and incubated at 37 °C with shaking at 180 rpm for an additional 5 hours. Following, two washing steps with fresh LB broth. B. subtilis cultures at an OD_600nm_ of 2.0 were mixed at a 1:4 ratio (prey: attackers). Five microliters of the mixture were spotted onto CM media for 40 hours at 30°C. Cells from the colonies were retrieved by resuspending in 2 mL of LB medium, and cellular viability per colony was estimated by serial dilution and plating on LB-agar supplemented with kanamycin (10 µg/mL) or spectinomycin (100 µg/mL). Each competition assay was performed independently in triplicate. For interbacterial competition monitored by time-lapse microscopy, after normalizing the optical density (OD) to 2.0, cells were mixed at a 1:1 ratio and pipetted onto a CM medium support, adapted for *B. subtilis* as described by ^44^.

### Bioinformatics

Structural predictions of T7bU and its interactions were conducted using sequences from *Bacillus subtilis* 168 (RefSeq: GCF_000009045.1), *Listeria monocytogenes* serotype 4b strain LL195 (RefSeq: GCF_000318055.1), *Clostridium acetobutylicum* ATCC 824 (RefSeq: GCF_000008765.1), and *Lactococcus cremoris* subsp. *cremoris* KW2 (RefSeq: GCF_000468955.1). The predictions were performed using UCSF ChimeraX (v1.4, 2022-06-03), which integrates the ColabFold-AlphaFold2 suite ^28,34,35^ and Alfafold3 server ^45^.

---

**Extended Data Figure 1.**
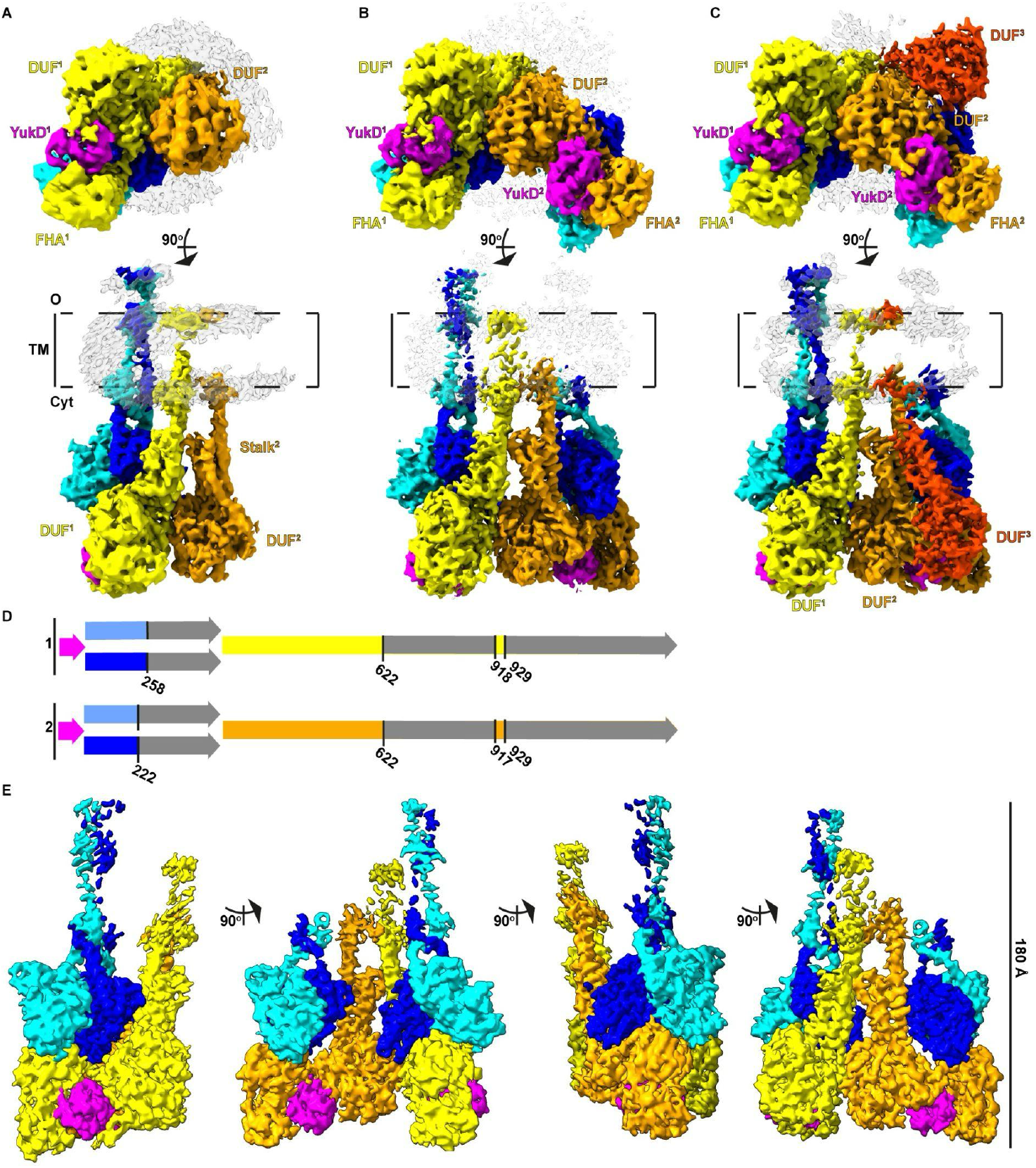
Cryo-EM Density Maps of T7bCU Multi-Complexes Revealing Assembly States. **(A–C)** Cryo-EM density maps showing three different degrees of oligomerization observed for T7bCU multi-complexes. Each state of oligomerization was achieved through single particle analysis (SPA). The maps in (A), (B), and (C) were derived from the highest-resolution structure (3.8 Å; 303,198 particles) presented in Figure 1A. **(A)** Monomeric T7bCU complex with an additional YukB DUF and stalk domain (134,631 particles, 4.2 Å). **(B)** Dimeric T7bCU complex (4.2 Å, 94,509 particles). **(C)** Dimeric T7bCU complex with an additional YukB DUF and stalk domains (4.2 Å, 73,157 particles,). YukB subunits 1, 2, and 3 of are colored yellow, orange, and red, respectively, with FHA, DUF and the Stalk domains. YukC subunits are colored blue and cyan, and YukD is shown in magenta. Key structural features, including transmembrane (TM) and cytoplasmic (Cyt) regions, are also annotated. Detergent belts are colored in semi-transparent grey. **(D)** Schematic representation of the density map of the dimer of T7bCU complexes. Electron densities observed are colored accordingly; gray areas represent regions not visible in the maps. **(E)** Cryo-EM density map of the dimer of T7bCU complexes, shown in 90° rotations. The models illustrate the organization of YukB, YukC, and YukD, as well as the oligomerization basis mediated by YukB subunits 1 and 2.

**Extended Data Figure 2.**
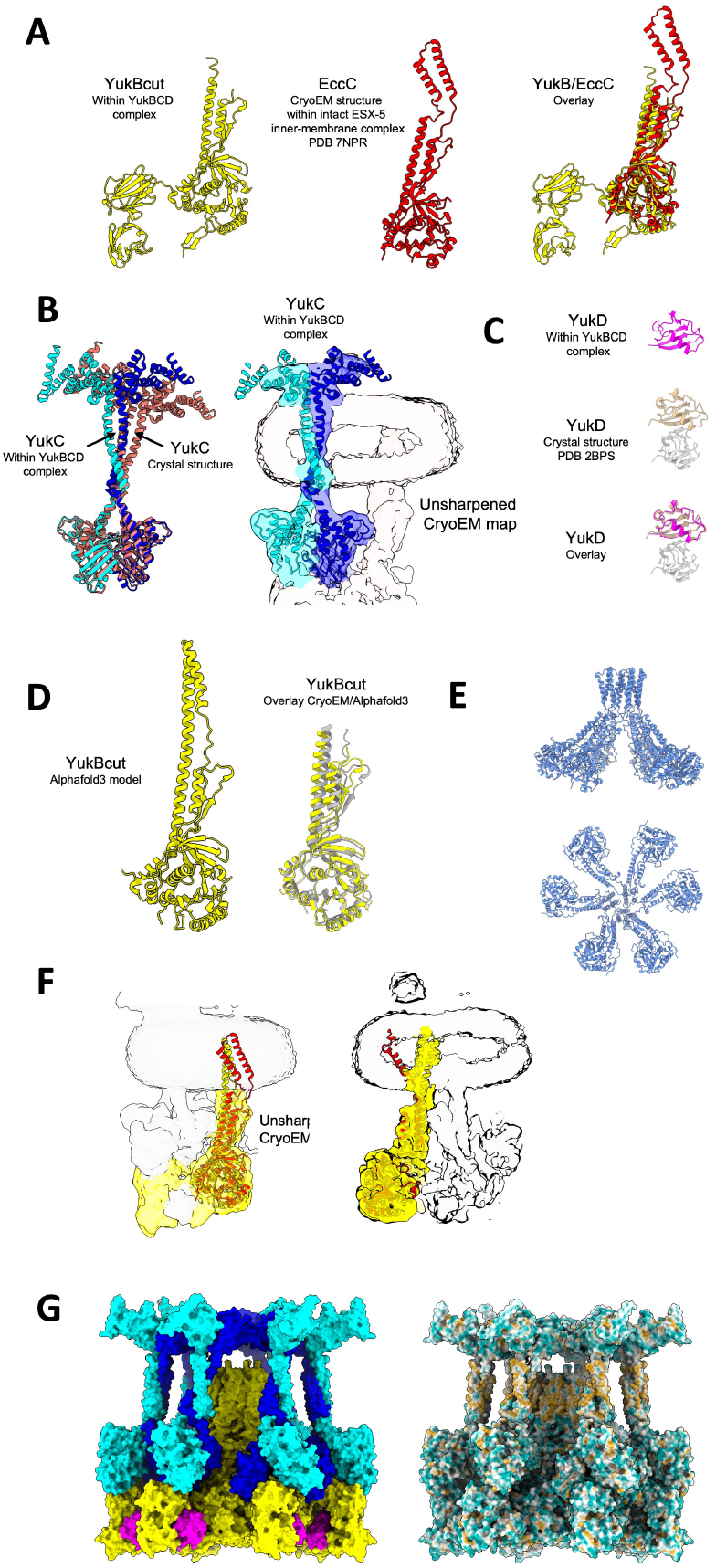
Structural comparisons between T7SSb components and their homologs. **(A)** Comparison between the YukB DUF and stalk domains with the corresponding regions in mycobacterial EccC (PDB: 7NPR), highlighting structural similarities. **(B)** Superposition of the YukC dimer from the CryoEM structure with the previously determined crystal structure (PDB: 6Z0F), showing the different angle of the transmembrane helices relative to the pseudokinase domain. Lower resolution densities enabled rigid-body docking of the transmembrane helices and C-terminal domains using the crystal structure as template. **(C)** Structural comparison between YukD from the CryoEM structure and its previously determined crystal structure (PDB: 2BPS), showing their high similarity. **(D)** Comparison between the DUF and stalk domains from YukB CryoEM structure and AlphaFold3 prediction. **(E)** CryoEM structure of the EccC hexamer showing the organization of DUF, Stalk and Transmembrane domains. **(F)** Comparison of transmembrane domain orientations between the YukB AlphaFold model and EccC CryoEM structure. The position of the TM domain in YukB is validated by the low resolution CryoEM map we obtained. **(G)** Surface representations of the composite model of the T7SSb hexamer showing YukBcut (Yellow), YukC (Cyan and blue) and YukD (magenta). Right panel shows surface representation according to hydrophobicity.

**Supplementary Figure 1.**
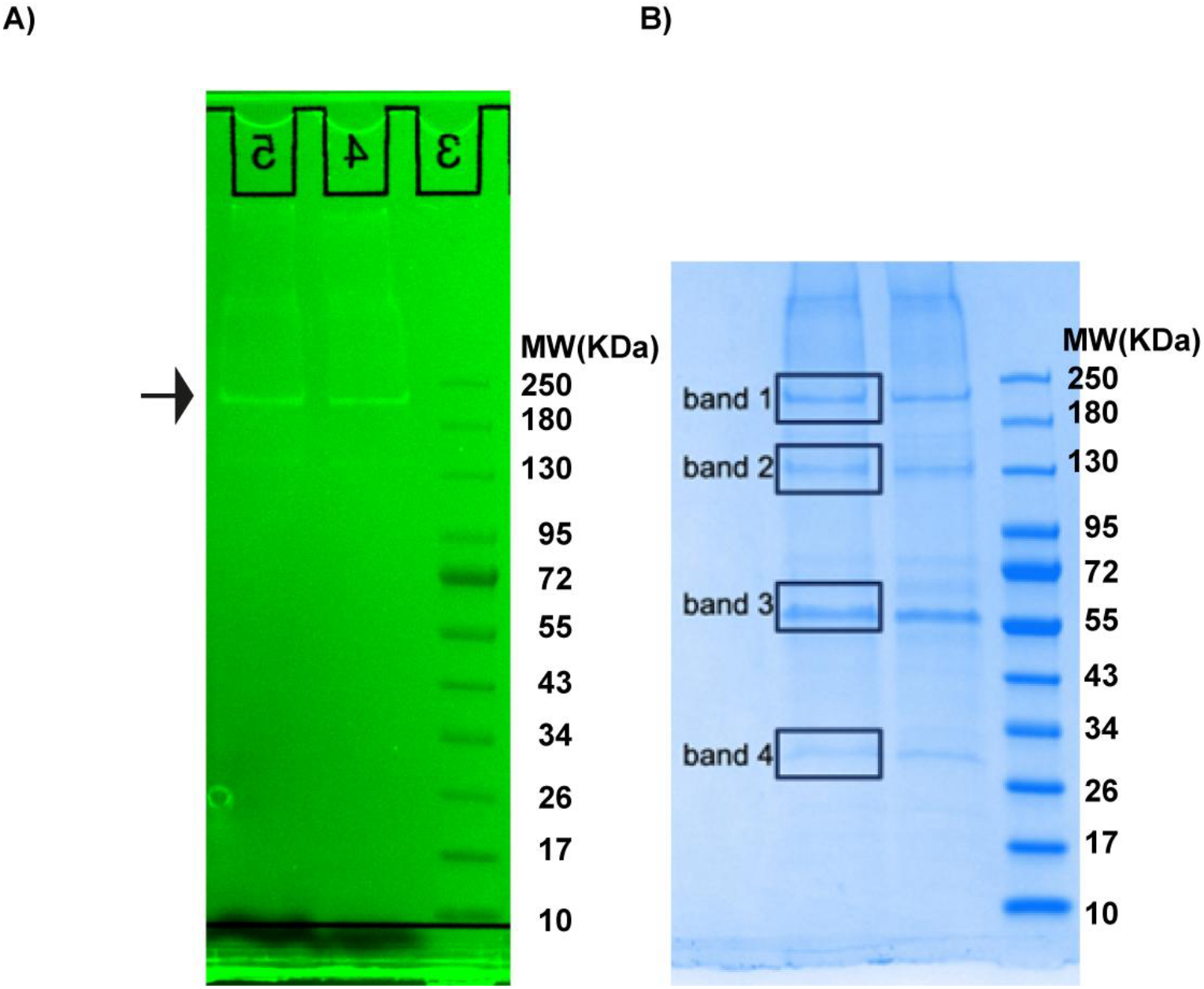
Purification of the T7SSb from *B. subtilis* expressing YukB-GFP-Twin-Strep tag (YukB-GFP-TST). SDS-PAGE showing the GFP fluorescence signal from the purification of YukB-GFP in *Bacillus subtilis*. The arrow indicates the fluorescent GFP signal, corresponding to a molecular weight of approximately ∼200 kDa from YukB-GFP-TST. A total of 50 µL of the concentrated detergent-solubilized eluted sample was obtained from the affinity chromatography (resin: Strep-Tactin® XT Sepharose), and 10 µL of this sample was applied to the gel in lane 4 and 5. GFP fluorescence was visualized using a Chem BioDoc imaging system (Bio-Rad). Molecular weight (MW) markers in Lane 3 (NEB: P7719S). **(B)** Coomassie-stained SDS-PAGE of the same gel shown in (A). Bands labeled 1, 2, 3, and 4 were excised and analyzed by mass spectrometry (MS). Band 1 was predominantly identified as YukB-GFP. A detailed list of the proteins detected in Bands 1 through 4 is provided below: **Band 1:** Main protein is YukB-GFP-TST. Other T7SS-related proteins found in the MS analysis : YukD (KEGG: BSU31900). **Band 2:** Main protein is Pyruvate Carboxylase (KEGG: BSU41860). Other T7SS-related proteins found in the MS analysis: YukB(KEGG: BSU31875), YukC(KEGG: BSU31890), YukD, YueB (KEGG: BSU31860). **Band 3:** Main protein is Ribose Import: ATP-binding protein RbsA (KEGG: BSU35940). Other T7SS-related proteins found in the MS analysis: YukC, YueB, YukD. **Band 4:** Main protein is Biotin Carboxyl Carrier Protein (BSU24350). Other T7SS-related proteins found in the MS analysis: YueC (KEGG: BSU31850), YukC, YueB, YukD, YueD (KEGG: BSU31840).

**Supplementary Figure 1.**
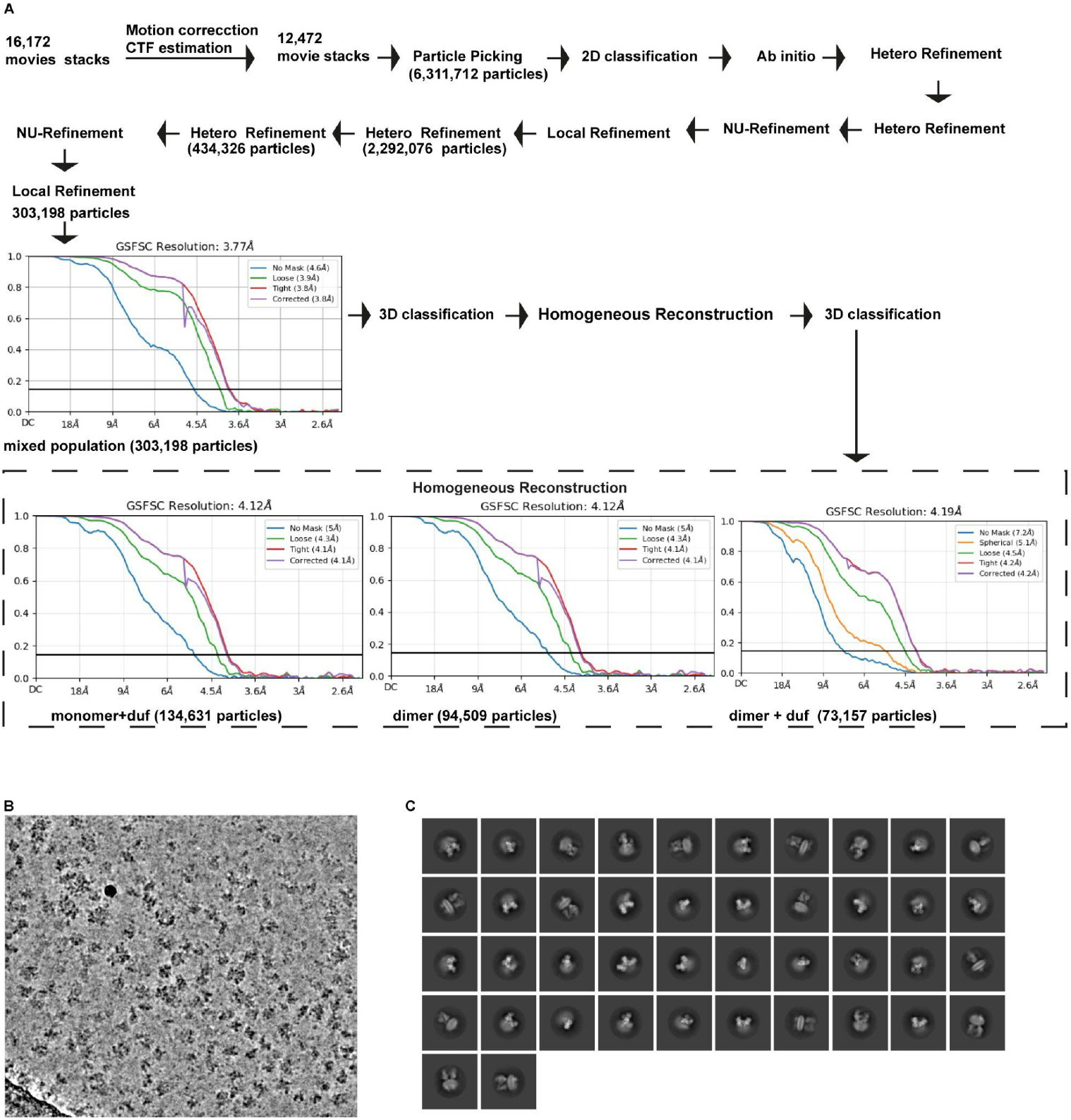
Single-Particle Analysis of the T7SSb Complex. **(A)** Cryo-EM data-processing workflow used to generate the final density map obtained in this study. Global Fourier Shell Correlation (GFSC) curves are shown for maps derived from focused refinement, containing a mixed population of monomer + DUF, dimer, and dimer + DUF particles, as well as the GFSC curves for the separated maps obtained after 3D classification and homogeneous refinement of the mixed population. **(B)** Example of a Cryo-EM micrograph from the dataset. **(C)** Representative 2D classes from the Cryo-EM data.

**Supplementary Figure 3.**
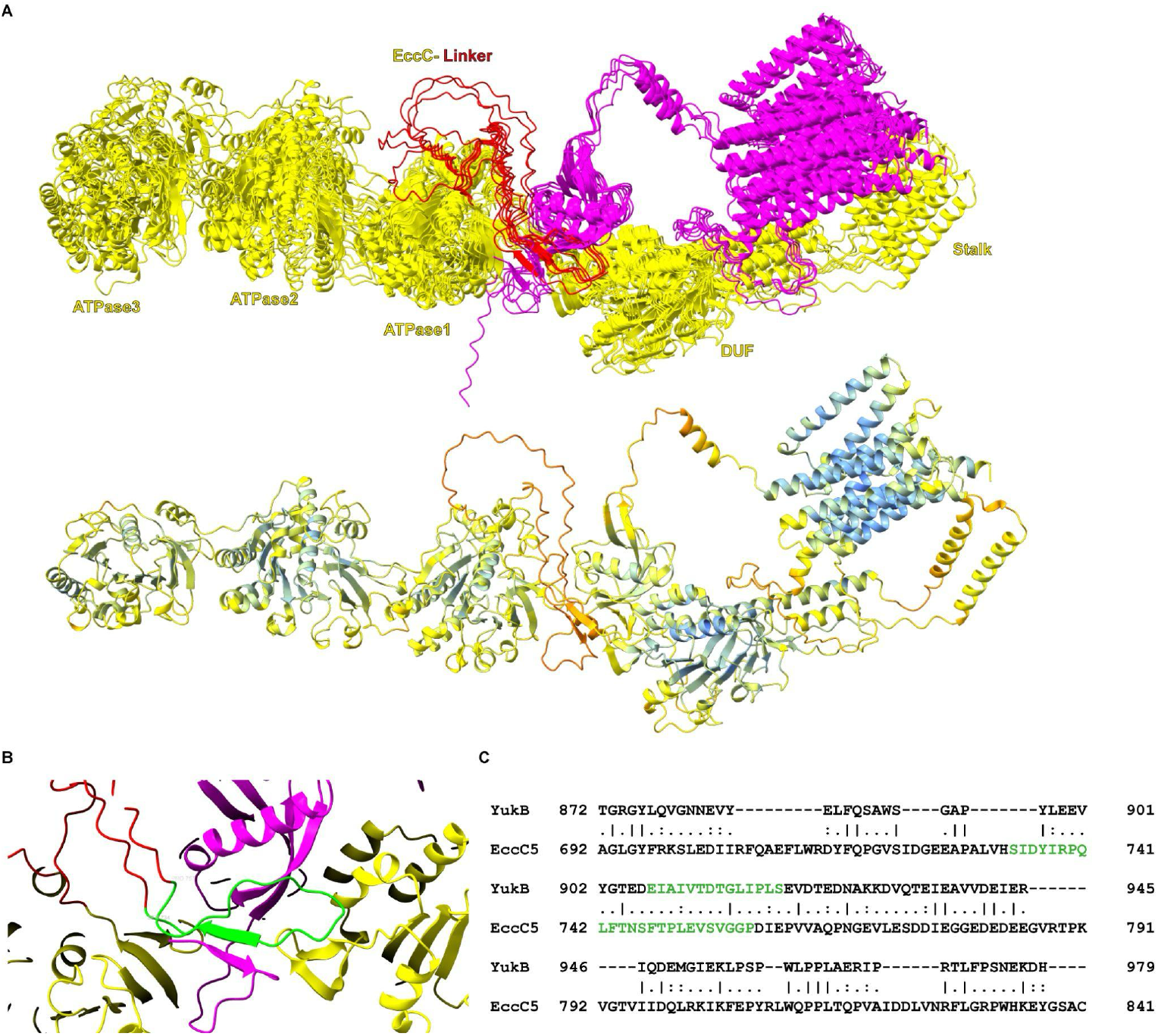
AlphaFold prediction of the EccC5-EccD5 complex from *Mycobacterium tuberculosis*. **(A)** Superimposition of five EccC5 models (yellow), showing ATPase1–3 domains and the stalk domain predicted by the AlphaFold3 Server^45^. The flexible linker (red), located between ATPase1 and ATPase2 (residues EccC5 711–791), suggests a potential interaction with the ubiquitin-like domain of EccD5 (magenta), consistent with the YukB-YukD interaction (Figures 1 and 2, results from this study). The lower panel is color-coded based on pLDDT confidence scores (blue for highest quality; yellow for low quality and orange for lowest quality). Sequences used for prediction: EccC5 (Uniprot: P9WNA5) and EccD5 (Uniprot: P9WNP90). For simplicity, the images show only a dimer of EccC5-EccD5. **(B)** Enlarged view showing a motif in EccC5 (residues 734–756, green) interacting with the N-terminal extension of EccC5 and the exposed beta strand of the ubiquitin-like domain of EccD5 (magenta). **(C)** Sequence alignment between EccC5 (Uniprot: P9WNA5) and YukB (Uniprot: A0A6M3ZG80) was performed using the EMBOSS Stretcher algorithm (https://www.ebi.ac.uk/Tools/psa/emboss_stretcher/) with default parameters and full-length sequences. The figure shows the aligned sequences, highlighting the flexible extension of EccC5, which includes the EccD5-interacting motif, in green. Similarly, the beta-hairpin region of YukB, containing the YukB-interacting motif, is highlighted in green.

